# Integrated single-cell multiome analysis reveals muscle fiber-type gene regulatory circuitry modulated by endurance exercise

**DOI:** 10.1101/2023.09.26.558914

**Authors:** Aliza B. Rubenstein, Gregory R. Smith, Zidong Zhang, Xi Chen, Toby L. Chambers, Frederique Ruf-Zamojski, Natalia Mendelev, Wan Sze Cheng, Michel Zamojski, Mary Anne S. Amper, Venugopalan D. Nair, Andrew R. Marderstein, Stephen B. Montgomery, Olga G. Troyanskaya, Elena Zaslavsky, Todd Trappe, Scott Trappe, Stuart C. Sealfon

**Affiliations:** Department of Neurology, Icahn School of Medicine at Mount Sinai (ISMMS), New York, NY 10029, USA; Lewis-Sigler Institute of Integrative Genomics, Princeton University, Princeton, NJ 08544, USA; Center for Computational Biology, Flatiron Institute, Simons Foundation, New York, NY 10010, USA; Human Performance Laboratory, Ball State University, Muncie, IN 47306, USA; Department of Medicine, Cedars-Sinai Medical Center, Los Angeles, CA 90048, USA; Department of Genetics, Stanford University, Stanford, CA 94305, USA; Department of Pathology, Stanford University, Stanford, CA 94305, USA; Department of Computer Science, Princeton University, Princeton, NJ 08544, USA

**Keywords:** acute endurance exercise, skeletal muscle, human vastus lateralis, single-nucleus multiome, snRNA-seq/snATAC-seq, circadian control, gene regulatory circuits, transcriptomics, epigenomics, PPAR𝜹

## Abstract

Endurance exercise is an important health modifier. We studied cell-type specific adaptations of human skeletal muscle to acute endurance exercise using single-nucleus (sn) multiome sequencing in human vastus lateralis samples collected before and 3.5 hours after 40 min exercise at 70% VO_2_max in four subjects, as well as in matched time of day samples from two supine resting circadian controls. High quality same-cell RNA-seq and ATAC-seq data were obtained from 37,154 nuclei comprising 14 cell types. Among muscle fiber types, both shared and fiber-type specific regulatory programs were identified. Single-cell circuit analysis identified distinct adaptations in fast, slow and intermediate fibers as well as *LUM*-expressing FAP cells, involving a total of 328 transcription factors (TFs) acting at altered accessibility sites regulating 2,025 genes. These data and circuit mapping provide single-cell insight into the processes underlying tissue and metabolic remodeling responses to exercise.

## Introduction

Physical activity provides diverse health benefits, including improvements in metabolism, immunity, cancer suppression, cardiac health and brain function.^1,2^ Skeletal muscle is directly perturbed by physical activity and is a primary mediator of these health benefits. Acute endurance exercise induces numerous physiological changes in skeletal muscle that initiate a complex regulatory program modulated at epigenetic and transcriptional levels.^3,4^ The molecular adaptations from repeated endurance exercise bouts lead to phenotypic changes within the muscle (i.e. improvements in aerobic fitness, exercise performance, and increase in lipid oxidation) as well as improved overall health.^5,6^

Skeletal muscle is a complex heterogeneous tissue composed of multinucleated fibers and mononuclear cells, such as immune cells, mesenchymal cells, and endothelial cells. Human skeletal muscle fibers include slow-twitch (slow) fibers, fast-twitch (fast) fibers, and intermediate fibers (*i.e.* fibers with both slow and fast characteristics). Each fiber type shows differing physiological responses to exercise that influence glucose metabolism, lactate production, and aerobic capacity.^3,7–17^ The skeletal muscle transcriptional program has been studied extensively at whole-tissue and pooled fiber type levels,^18–23^ Because of the importance of the cross-talk between skeletal muscle and diverse organs in modifying health and disease, an ongoing National Institutes of Health program (Molecular Transducers of Physical Activity Consortium, MoTrPAC) is developing a comprehensive whole-tissue molecular map of response to exercise.^24^

Given the heterogeneous cellular composition of muscle, it is necessary to resolve the regulatory landscape at a single-cell (sc) level. However, no studies to date have examined exercise response in complete human skeletal muscle tissue at sc resolution. Several studies have performed scRNA-seq on human skeletal muscle mononuclear cells at baseline, although most have focused on muscle stem cells.^25–30^ Lovrić et al. analyzed the exercise response of mononuclear skeletal muscle cells via scRNA-seq.^31^ Additionally, although multiple studies have performed bulk tissue ATAC-seq on skeletal muscle at baseline,^32–37^ no studies have characterized the changes in chromatin architecture in response to endurance exercise in bulk or single-cell human skeletal muscle tissue. Such a study is needed to provide insight into the regulatory mechanisms underlying the gene response to exercise and to improve the interpretation of the extensive and ongoing exercise research generating whole tissue muscle data.

A major step towards elucidating the role of muscle cell subtypes in mediating the systemic effects of exercise is to identify the regulatory circuitry (comprising the gene, the transcription factor (TF) regulating the gene, and the specific upstream interaction site between the TF and the gene) that underlies the control of gene expression within individual cells. SC multiome assays, which simultaneously determine gene expression and chromatin accessibility levels within the same cells, have the potential to both resolve the molecular changes in response to exercise and to allow the underlying sc gene regulatory circuitry to be reconstructed. By analyzing sc multiome data from subjects at baseline and after an acute bout of endurance exercise, we map the regulatory landscape within individual cell types. Our study includes non-exercising control subjects that are sampled at the same time of day as the exercise subjects, allowing us to differentiate exercise responses from circadian rhythm effects. Our results provide a comprehensive single-cell integrated skeletal muscle atlas of epigenomic and transcriptomic exercise responses and their underlying molecular circuitry.

## Results

### Same-cell single-nuclei multiome sequencing provides cell-type resolution map of exercise response

In order to generate a multiome muscle exercise dataset, vastus lateralis skeletal muscle biopsies were obtained from four subjects (two males, two females) before and 3.5 h after an acute endurance exercise bout. To control for circadian effects, one male and one female resting control subject were sampled and analyzed at the same time of day as the exercised subjects (Figure 1A, Table S1). Nuclei isolated from the twelve skeletal muscle samples underwent single-nuclei (sn) multiome library construction and sequencing to generate same-nucleus RNA-seq and ATAC-seq datasets. The 37,154 cells from all samples passing quality control (QC) for both RNA-seq and ATAC-seq were integrated, clustered, and annotated into 14 cell types using previously identified cell type markers^30^ (see Methods) (Figure 1B). The main markers used to annotate cell types are shown in Figure S1. Cell-type proportions were largely consistent across groups, sexes, and time-points (Figure 1C, Figure S2). Natural Killer (NK) cells showed a trend towards an increase from pre-to post-exercise consistent with a previous report,^31^ but this increase did not achieve significance after adjustment for multiple hypothesis testing (Figure S2A).

**Figure 1:**
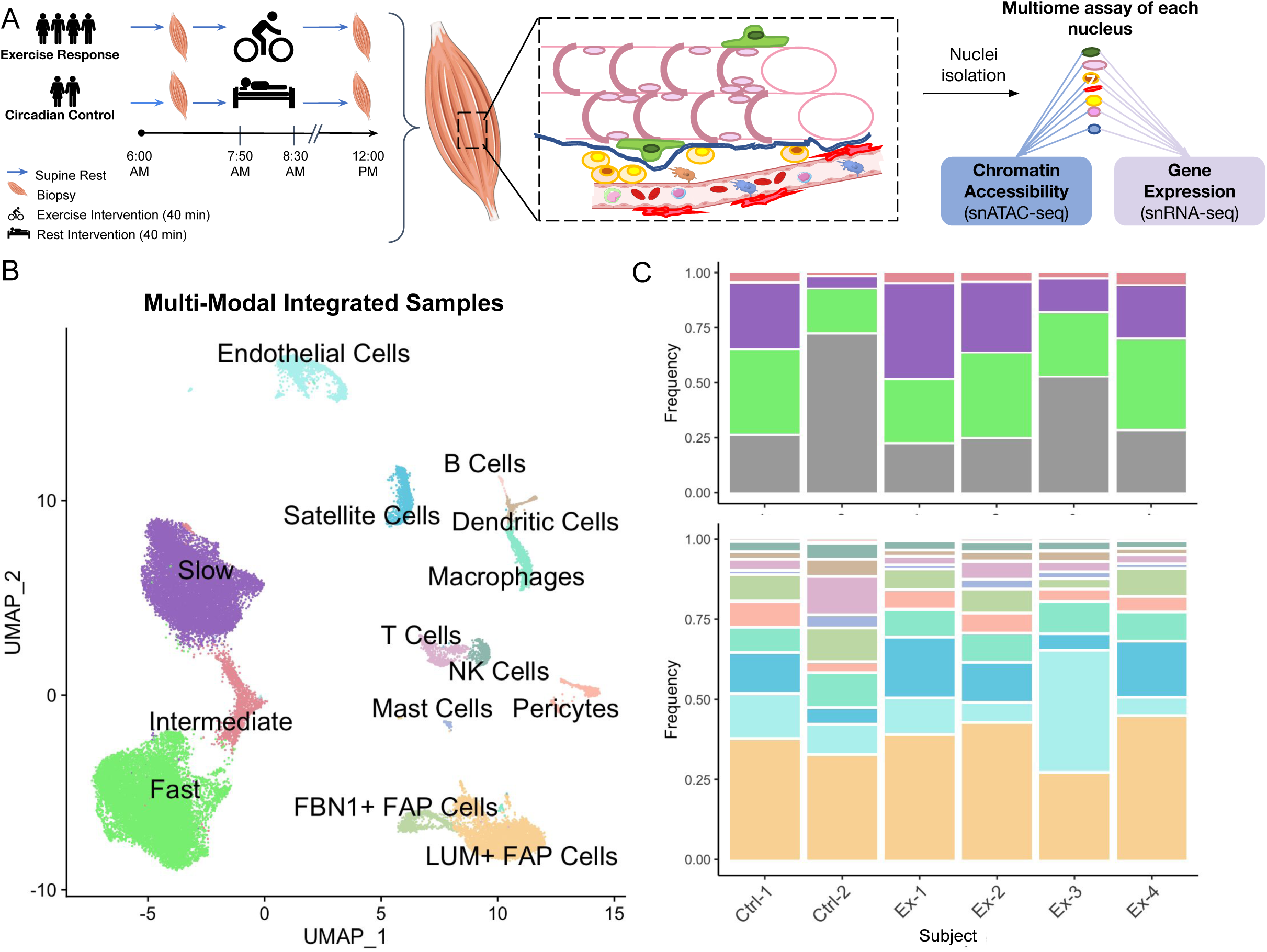
Generation of cell-type resolution map of exercise response. (A) Human vastus lateralis biopsies were collected before and 3.5 hours post 40 min exercise at 70% VO_2_max in four subjects as well as biopsies at the same time-points from two supine resting circadian controls. Nuclei were isolated and underwent same-cell single-nucleus multiome assay to assess gene expression and chromatin accessibility simultaneously. (B) High quality single-nuclei assay data was integrated and clustered with fourteen cell-types identified. See Figure S1 for cell-type markers used for annotation. (C) The proportion of each cell-type from the single-cell analysis within each baseline subject is shown. Plot is split by all cell-types (top panel) and mononuclear cells only (bottom panel) to enable easier comparison of low-frequency cell-types. Total frequency of mononuclear cells is shown in gray in top panel. Column labels denote group identity (Ctrl: circadian control; Ex: exercise response) and subject ID. Subjects Ctrl-1, Ex-1, and Ex-2 are female; subjects Ctrl-2, Ex-3, and Ex-4 are male. See Figure S2 for a detailed examination of cell-type proportions.

To investigate molecular changes within cell types due to exercise, we first identified the pre-to post-exercise differentially expressed genes (DEG) and differentially accessible regions (DAR), excluding features showing comparable circadian responses (see Methods). As an example of the importance of considering circadian regulation, *FKBP5* is downregulated in both the circadian control group and the exercise response group (*P* < 0.001 and log_2_ fold change < -2.0) (Figure S3A,B). However, our results suggest that the regulation of *FKBP5* is primarily due to circadian effects. Over all cell types, changes observed in 1,210 unique genes and 1,068 unique chromatin regions were attributed to circadian regulation effects (Figure S3C,D) and were excluded from further analysis of exercise-related responses. Gene or chromatin responses regulated by circadian effects that could be identified as showing a synergistic effect of exercise were retained.

Fast fibers, slow fibers, and LUM+ FAP cells had the greatest numbers of exercise-regulated DEGs and DARs (Figure 2A,B). While the numbers of fast and slow fibers in these analyses were comparable (n = 13,564 fast fibers and n = 10,444 slow fibers), there were more differentially expressed transcripts in fast fibers and nearly double the number of significantly regulated chromatin accessibility sites. Fast and slow fibers showed both overlapping and fiber-specific regulated genes and chromatin regions (Figure 2C, S3E,F). A total of 596/2281 (26%) DEGs found in any cell type in our analysis were previously identified as exercise-regulated in a skeletal muscle bulk transcriptome meta-analysis of 15 studies (Figure 2C).^23^

**Figure 2:**
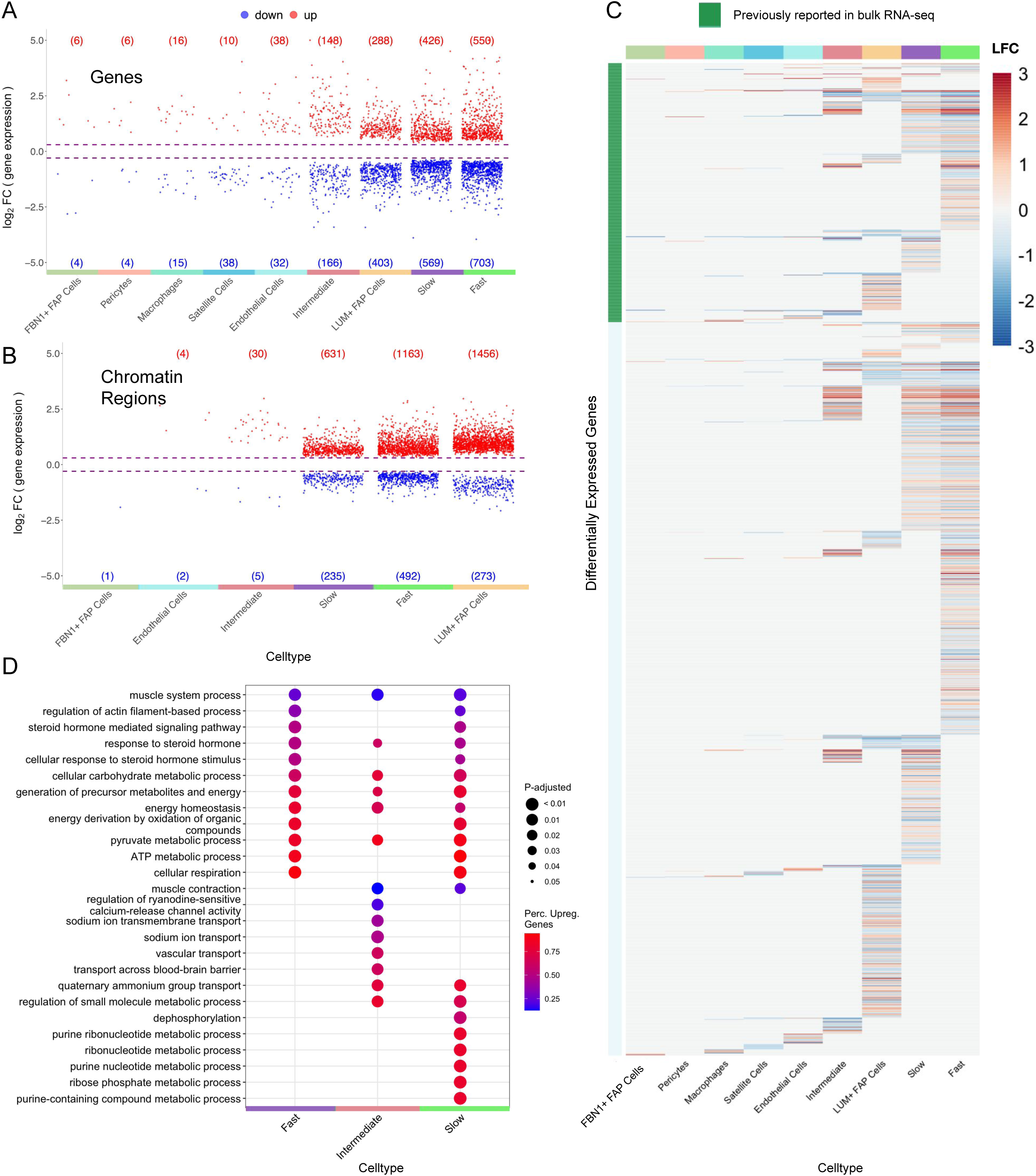
Cell-type resolution map of exercise response enables differential analysis which reveals high differential expression and accessibility for fiber types and LUM+ FAP cells. (A-B) Volcano plots show the distributions of upregulated and downregulated genes (A) and chromatin regions (B) for each cell-type. Each point denotes a differentially expressed gene (A) or differentially accessible region (B) for a given cell-type. Cell-types with no significant differential features are not shown. Circadian-regulated features are excluded and shown in Figure S3C,D. (C) Heatmap shows overlap in differentially expressed genes within cell-types. Annotation bar denotes whether a gene has previously been validated as differentially expressed in bulk RNA-seq meta-analysis study (green). Color represents log_2_ fold change of each gene for each cell-type. (D) Enrichment analysis of biological processes in fiber types shows functionally different transcriptome profiles for each cell-type. Color represents the percentage of upregulated genes in a given pathway and larger dot radius denotes more significant biological function. See Figure S4 for enrichment analysis for all cell-types. See methods for detailed description of differential analysis including statistical tests used.

We performed enrichment analysis on the exercise-response genes to identify pathways that are most upregulated and downregulated for each cell-type (Figure 2D, Figure S4). Slow, fast, and intermediate fiber types shared multiple energy production/cellular respiration, steroid hormone response, and muscle function processes. Additionally, several molecular transport pathways were uniquely regulated in intermediate fibers and many ribonucleotide/nucleotide metabolic pathways were uniquely regulated in slow fibers.

Exercise stimulates remodeling of the muscle structure. Both satellite cells and LUM+ FAP cells had upregulated pathways for axon guidance. Satellite cells, LUM+ FAP cells, fast fibers, and slow muscle fibers were enriched in muscle remodeling and morphogenesis-related pathways. Endothelial cells showed exercise-activation of angiogenesis and wound healing pathways.

Of note, circadian rhythm pathways were downregulated by FBN1+ FAP cells and satellite cells in response to exercise. This is due to further regulation by exercise of some circadian-effect genes (e.g. *per1* and *per3*), highlighting the importance of including circadian control subjects in this study.

### PLIER analysis identifies conserved training responses across both gene expression and chromatin accessibility data in slow, fast, and intermediate fiber types

We used the Pathway Level Information Extractor framework (PLIER)^38^ to identify latent variables (LVs) which provide single value estimates of groups of genes and chromatin loci showing correlated patterns of activity within each cell type across samples. We analyzed the gene expression from RNA-seq and gene promoter accessibility from the ATAC-seq data, for each of the three muscle fiber types (see Methods, Figure S5A).

Among the gene-chromatin LVs identified by PLIER, two were regulated by exercise (Figure 3). One showed increases with exercise at the individual gene expression level and the corresponding promoter accessibility (Figure 3A). These changes were significant in all three fiber types for gene expression and in fast and slow fibers for chromatin accessibility (Figure 3B). The features of this upregulated LV were annotated to TCA cycle, pyruvate metabolism, and lysosome pathways (Figure S5B). In contrast, an LV associated with mTOR, spliceosome, and circadian expression pathways showed significant decreases in gene expression and chromatin accessibility in all three fiber types (Figure 3C-D, Figure S5B). In both of these exercise-response LVs, there was no significant change in circadian control subjects (Figure 3B,D).

**Figure 3:**
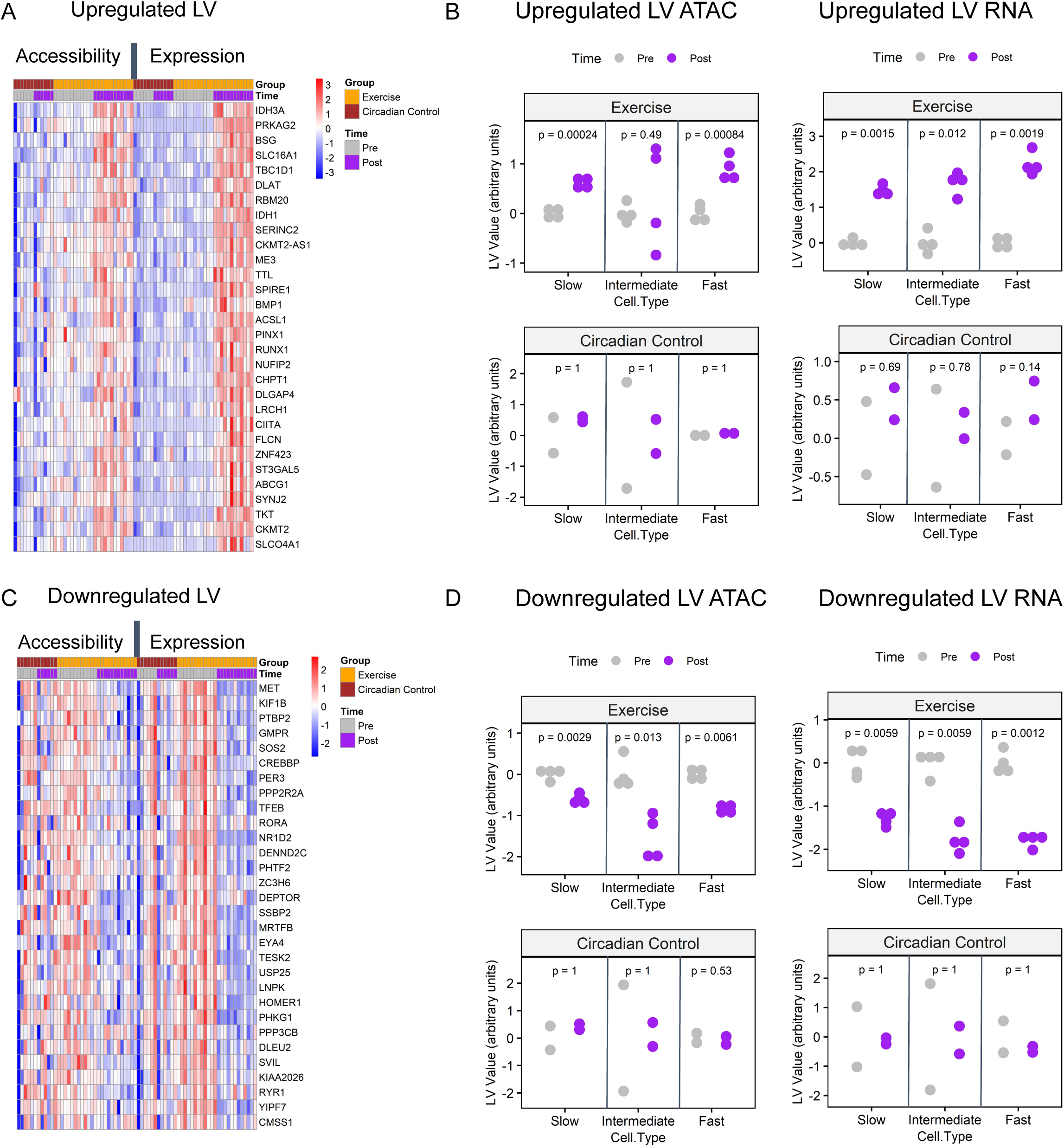
Latent variable analysis identifies conserved training responses across both RNA-seq and ATAC-seq data in slow, fast, and intermediate fiber types. Two latent variables (LVs) representing patterns of upregulation, LV9, (A,B) and downregulation, LV12 (C,D) conserved across both gene expression and promoter chromatin accessibility are shown. For complete PLIER latent variable analysis, see Figure S5. (A,C) Heatmaps represent chromatin accessibility and gene expression z-scored pseudobulk values for the top thirty genes that are associated with each LV. Each column represents one fiber-type sample; columns are ordered by ome type, exercise group, and time-point. (B,D) Scatterplots represent the summary values for exercise-regulated LVs. Plots are faceted by ome type (RNA vs. ATAC), exercise group (exercise vs. circadian control), and fiber type. Change between pre and post LV summary values is assessed via Holm-Bonferroni adjusted paired t-test with ns: p > 0.05, *: p <= 0.05, **: p <= 0.01, ***: p <= 0.001, ****: p <= 0.0001.

### Regulatory circuitry of exercise reveals molecular mechanisms underlying exercise adaptation

*Cis*-gene regulatory circuits, comprising genes, transcription factors (TFs) and their specific *cis*-regulatory sites in the chromatin, serve a major role in determining gene expression. The sc multiome datasets, by profiling the regulatory circuit components within each nucleus, make it possible to reconstruct the cell type-specific gene control circuitry mediating the transcriptional response to exercise at single-cell resolution (Figure 4A). Exercise-regulated circuits were identified for each cell type with concordant changes in target gene expression, TF expression and chromatin accessibility of regulatory sites between pre- and post-exercise samples (see Methods). For these analyses, we focused on fast, intermediate, and slow fibers, as well as LUM+ FAP cells, as these cell types had adequate cell numbers and robust responses to exercise for regulatory circuit inference. Applying recently developed complementary bioinformatic methods,^39,40^ a total of 2,025 exercise-regulated circuit genes were identified among the four cell types, with the highest number found in the fast fibers (Figure 4B). While some circuits were shared among cell types, the slow and fast fiber types each displayed distinct regulatory programs regulating hundreds of unique genes reflecting fiber and cell type-specific transcriptional remodeling programs activated by acute endurance exercise (Figure 4B). Interestingly, the regulatory programs showed a higher level of overlap among fiber types at the TF level in comparison with the regulated circuit genes (Figure 4C).

**Figure 4:**
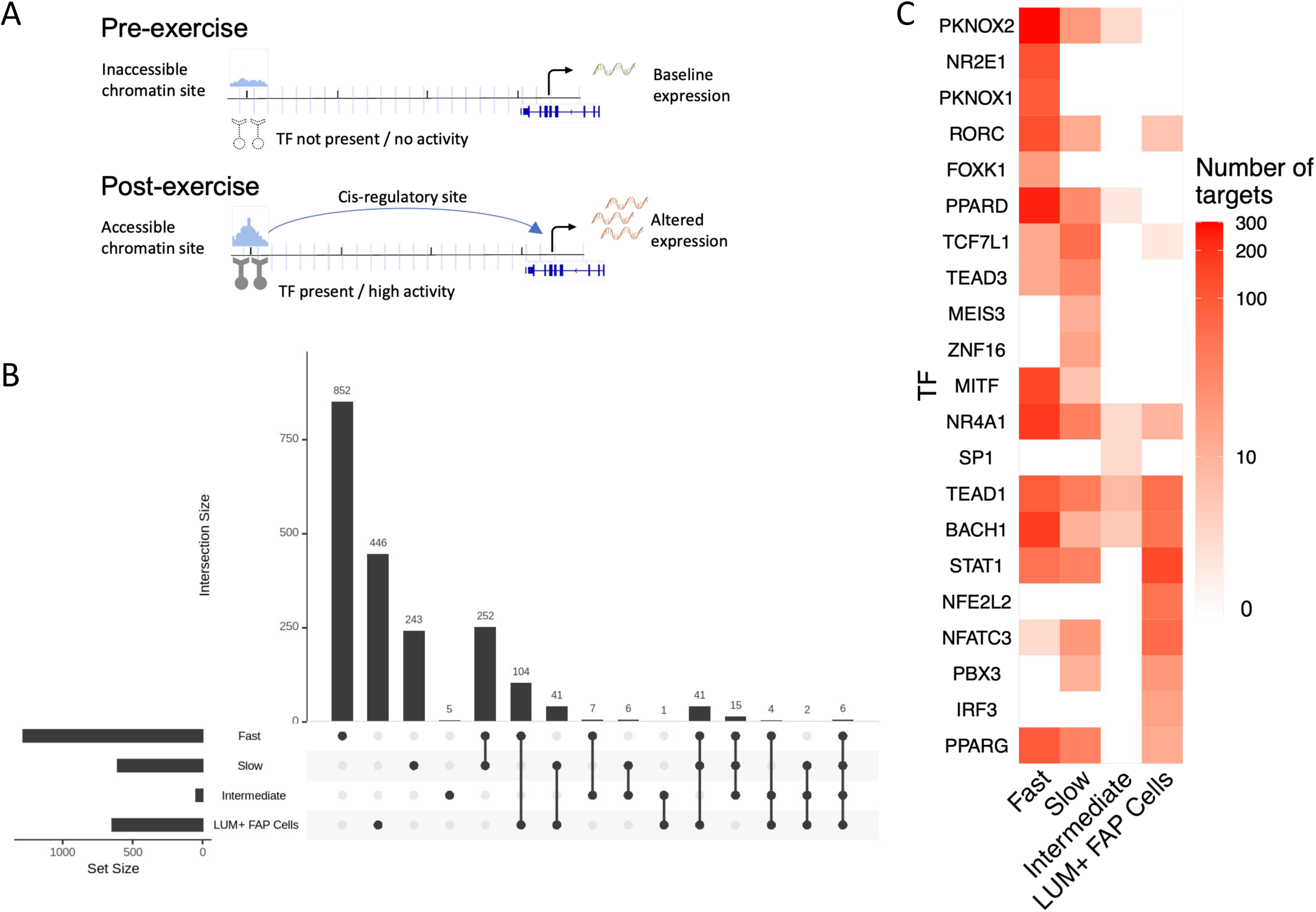
Regulatory circuitry of exercise reveals molecular mechanisms underlying exercise adaptation (A) Schematic figure showing the components of regulatory circuits underlying exercise-related gene expression changes. (B) UpSet plot displays the sizes of overlapping and unique sets of exercise-related circuit-genes identified in each cell-type. (C) Heatmap reveals the numbers of identified targets of selected TFs (rows) in each cell-type (columns).

To demonstrate the utility of the exercise sc multiome and inferred regulatory circuitry resource for understanding the mechanisms used by muscle to remodel metabolic processes, we focused on the TF PPAR𝜹 which has been implicated in regulation of glucose uptake, lipid oxidation, and muscle fiber switching.^41,42,43^ An example of a PPAR𝜹 regulatory circuit, in which a target gene *FABP3* was found to be regulated with exercise in fast fibers, is shown in Figure 5A. In this circuit, the TF (PPAR𝜹) and its target gene (*FABP3*) were both upregulated in response to exercise, while the PPAR𝜹 binding site upstream of *FABP3* also showed higher accessibility post exercise.

**Figure 5:**
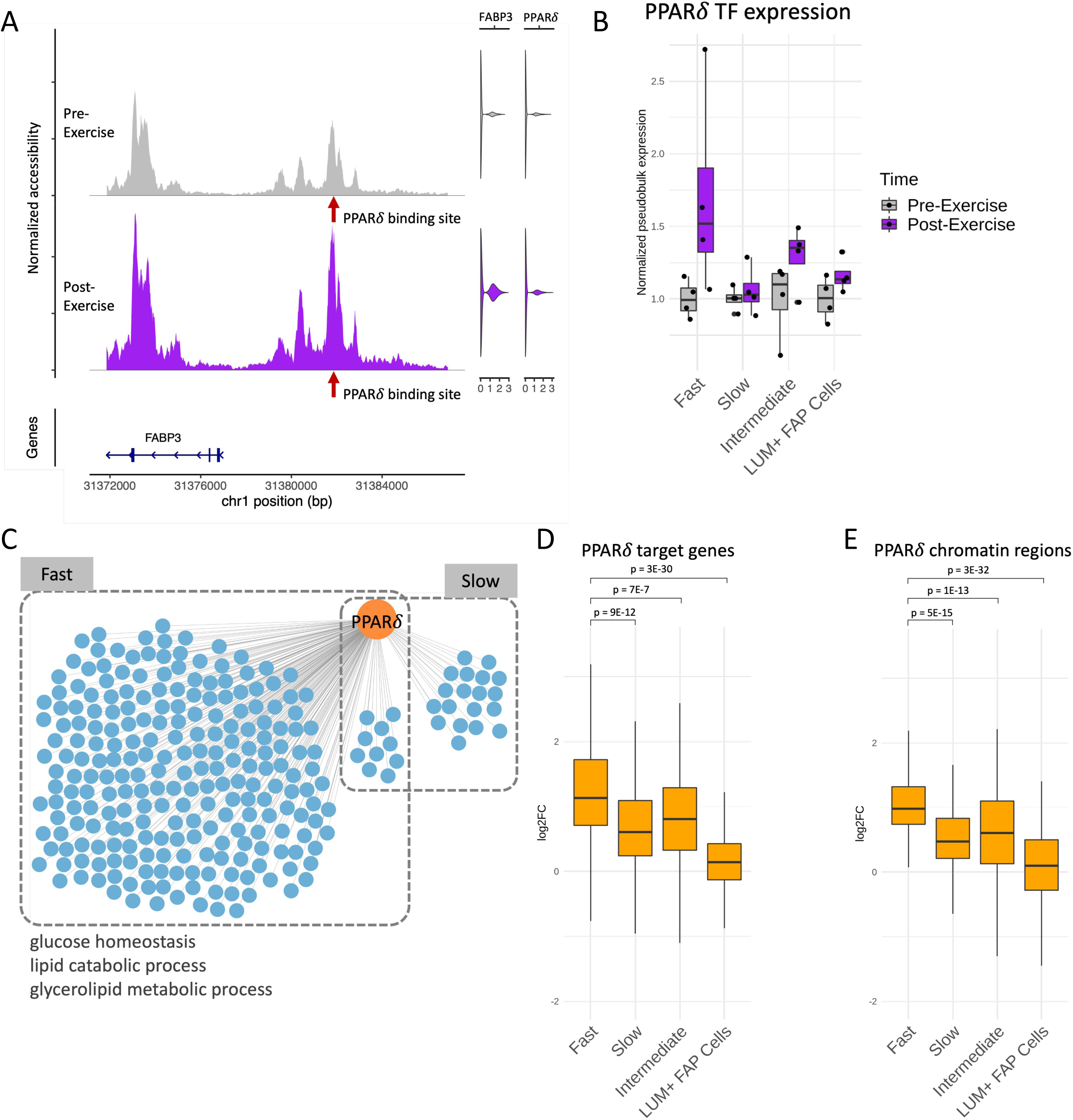
Regulatory circuitry of a fast fiber specific exercise-related TF: PPAR𝜹. (A) Example of a regulatory circuit (PPAR𝜹-*FABP3*) where TF PPAR𝜹 interacts with a cis-regulatory region upstream of the target gene *FABP3* and regulates its expression after exercise. Left panel shows the normalized chromatin accessibility of the genomic regions around the transcription start site (TSS) of *FABP3* in fast fibers pre-exercise (gray) and post-exercise (purple). Red arrow indicates the location of the PPAR𝜹 binding site. Right panels are violin plots showing the expression of *FABP3* and PPAR𝜹 pre-exercise (gray) and post-exercise (purple). (B) Normalized pseudobulk RNA expression of PPAR𝜹 in each group, time-point and cell-type is depicted as a boxplot with one point per sample. For each group and cell-type, both the pre-exercise and post-exercise RNA exercise of each sample were normalized by the mean of the pre-exercise samples. (C) Network plot reveals the cell-type specific target genes (blue) of PPAR𝜹 (orange) identified by the integrated regulatory circuit analysis. Selected GO terms enriched in the target genes are annotated below. (D-E) Log_2_ fold change (log_2_FC) between post-exercise and pre-exercise of the target genes (D) and chromatin regions (E) in the fast fiber specific PPAR𝜹 circuitry in each cell-type. P-values are calculated using a one-sided Wilcoxon test comparing the log_2_FC between fast fiber and other cell-types.

We next investigated the broader PPAR𝜹 regulatory network in multiple cell types, examining the three aspects of PPAR𝜹 circuits: PPAR𝜹 itself, circuit target genes, and associated chromatin regions. The RNA-level expression of PPAR𝜹 showed upregulation specifically in the fast fiber type in response to exercise (Figure 5B). PPAR𝜹 was also found to be a regulator of 242 circuit genes in fast fibers and only 29 circuit genes in slow fibers (Figure 5C, Table S4). To explore the extent of the PPAR𝜹 circuits’ specificity to fast fibers, we examined the differential signal of the fast fiber PPAR𝜹 circuit components across cell types. These 242 PPAR𝜹 circuit genes and 198 associated chromatin regions containing the PPAR𝜹 binding sites showed significantly higher upregulation in the fast fibers compared to other cell types (Figure 5D). These results suggest that PPAR𝜹 predominantly regulates a fast fiber specific network in response to endurance exercise.

Enrichment analysis of fast fiber PPAR𝜹 circuit genes revealed that metabolic pathways such as glucose homeostasis and lipid catabolic process were upregulated in the PPAR𝜹 network (Figure 5C), which is consistent with previous reports of PPAR𝜹’s functions in muscle fibers^41,42^ as well as the enriched GO terms in the standard functional enrichment analyses shown in Figure 2C. The discovery of the fine-grained regulatory circuits enabled us to identify the mechanisms behind the regulation of these important metabolic functions specifically within the fast fiber.

## Discussion

Muscle is a key tissue in mediating the metabolic changes occurring with endurance exercise that underlie many of its systemic effects on physiology and health. ^1^ We profiled the regulatory landscape of the response to endurance exercise in skeletal muscle using same-cell sc multiome data. The application of recently developed bioinformatic frameworks generated high-resolution reconstruction of the exercise-response gene regulatory circuitry at a cell-type specific level. The inclusion of resting control subjects allowed exercise responses to be distinguished from concomitant circadian changes.

Our analyses identified a total of 2,025 exercise-regulated circuit genes, as well as the TFs and interaction sites mediating the regulation of each gene. As we demonstrate with a detailed analysis of the PPAR𝜹 circuitry, our data and analyses provide deep insight into the gene regulatory mechanisms controlling any specific gene response within muscle cell types.

We have recently determined using a rat animal model that the transcriptome response to endurance exercise training is largely distinct among different tissues.^44^ Notably, we now find in human muscle that the transcriptome response to acute endurance exercise is largely distinct among the different cell types within the same tissue. Further, we find that each cell type, including the slow, fast, and intermediate muscle fiber types, show largely distinct responses to exercise, with fast fibers showing the broadest and most robust activation of transcriptional and chromatin remodeling programs. Interestingly, fast and slow fibers had a high degree of overlap in circuit transcription factors and a lower degree of overlap in circuit genes, suggesting that a major mechanism contributing to fiber-type specific responses to exercise may be the capacity of genes to respond in each tissue due to their baseline epigenetic state.

Our results help elucidate the mechanisms of exercise-mediated metabolic changes at the cell-type level. Pathway enrichment analysis of fiber-type differential genes revealed that several pathways involving energy production and cellular respiration were upregulated in all fibers with greater regulatory changes seen in fast fibers. The coordinated upregulation of genes and chromatin sites concentrated on metabolic pathways present in all muscle fibers, but of greater amplitude in fast fibers, was also detected by both gene pathway analyses and by integrated gene-chromatin latent variable deconvolution (Figure 3). Notably, purine metabolic pathway regulation related to the need for increased energy production was concentrated in slow fibers. Purine metabolism is commonly considered to be increased with higher intensity anaerobic exercise or exercising in an energy depleted state.^45–47^ While slow fibers should be heavily activated during the 40 minutes of cycle exercise at ∼70% of VO_2_max completed by these subjects, the energy demands of both fiber types would be expected to be met almost exclusively with aerobic energy sources after the first few minutes of the exercise bout.^48,49^ Also, given the pre-trial controls, subject fitness status, and duration of the exercise bout, we would not expect intramuscular ATP levels or energy stores (*i.e.*, glycogen) to be very low in any fiber type during this exercise bout.^48,49^ Our results indicate the purine transcriptional pathway is still activated in slow fibers by acute aerobic exercise. The slow fiber-type predominance is consonant with findings that slow muscle has a greater capacity for purine metabolism in animals.^46^ Overall, the current findings suggest a potential novel role of altered human skeletal muscle purine pathway energy production initiated by endurance exercise in a muscle fiber-type specific manner that warrants further study.^50^

PPAR𝜹 is an important factor in glucose homeostasis and muscle substrate regulation. Our analyses now resolve the relative role of distinct fiber types in this process and the specific PPAR𝜹 regulatory circuits responding to exercise within each fiber type. We find that PPAR𝜹 primarily regulates a fast fiber specific network in response to endurance exercise. This is interesting given the centrality of PPAR𝜹 for the shift in substrate utilization from glucose towards fat associated with endurance exercise training.^51^ These results also demonstrate how the data and analysis resource we have generated provides detailed exercise-response cell and circuit level resolution for any transcription factor or gene of interest.

We demonstrate the value of considering circadian effects in exercise response genomic studies. Our high-resolution transcriptome regulation data identify many of the regulated genes reported in a meta-analysis of 309 bulk transcriptome samples. However, while 596 genes we identified were previously detected by bulk analysis in that study, 283 genes previously reported to be exercise-related are attributed by our analysis to circadian effects. Thus, in addition to providing the first exercise-response sc multiome or chromatin accessibility data, our study distinguishes the transcriptome and chromatin remodeling effects due to exercise from predominantly circadian related changes (Figure S4).

Our resource of within-single-cell gene expression and chromatin accessibility of the effects of endurance exercise and cell-type resolution map of the gene regulatory circuitry underlying the effects of exercise is made available to the community in a web accessible interface. Given the prominent role skeletal muscle plays in locomotion, overall metabolic health, and organ cross-talk, these results provide a framework for future studies into the mechanisms through which the exercise effects on muscle modulate health and disease.

### Limitations of Study

Young healthy adults may not reflect changes across lifespan. Additionally, we lack power for accurately studying sex differences. We are studying exercise response at single-nucleus resolution, which does not precisely reflect the multinucleated structure of muscle fibers.

Integrated studies of spatial transcriptomics and chromatin structure multiome methods, which have recently been developed,^52^ may be helpful in resolving the relationship of regulatory changes of nuclei within the same individual fibers. The timing of post-exercise sample collection was limited to a single time point (3.5h post-exercise) and more information is likely to be gained by additional time-points, both earlier and later than the current post-exercise time-point. Furthermore, the study is limited to endurance exercise at one level of intensity; results may differ depending on exercise modality and intensity.

#### Data and Code Availability

Single-cell multiome data are available at the GEO repository under accession number GSE24006.

In addition, within-single-cell gene expression and chromatin accessibility of the effects of endurance exercise as well as a cell-type resolution map of the gene regulatory circuitry underlying the effects of exercise is available in a web accessible interface located at https://rstudio-connect.hpc.mssm.edu/muscle-multiome/.

## Acknowledgments

This work was supported by funding from NIH grants R01MH125244, as well as funding via institutional Icahn School of Medicine at Mount Sinai funds and the Human Bioenergetics Program at Ball State University. We acknowledge the New York Genome Center for sequencing. This work was supported in part through the computational resources and staff expertise provided by Scientific Computing at the Icahn School of Medicine at Mount Sinai and supported by the Clinical and Translational Science Awards (CTSA) grant UL1TR004419 from the National Center for Advancing Translational Sciences.

## Author Contributions

Conceptualization: T.T., S.T., S.C.S.; resources: T.T., S.T., T.L.C.; investigation: T.T., S.T., T.L.C., F.R.Z., N.M., V.D.N., M.S.A.; data curation: A.B.R, W.C., M.Z.; formal analysis: A.B.R., X.C., Z.Z., G.R.S., E.Z., W.C., O.G.T., A.R.M., S.B.M.; visualization: A.B.R., G.R.S., X.C., Z.Z., F.R.Z., S.C.S, E.Z., O.G.T.; writing − original draft: A.B.R., E.Z., S.C.S., X.C., Z.Z., G.R.S., F.R.Z., T.L.C.; writing – review and editing: all authors.

## Methods

### Human participants and muscle sample collection

Six participants were enrolled in the study, of which four subjects (2 males, 2 females) were in the endurance exercise group and two subjects (1 male, 1 female) were in the circadian control group. Table S1 includes information on subject sex, age, height, weight, BMI, body fat percentage, VO2max (absolute and relative), ethnicity, and group (circadian control vs. endurance exercise). Skeletal muscle biopsies were obtained from the vastus lateralis following local anesthetic (1% lidocaine HCl) with a 6-mm Bergström needle with suction^18,53,54^ both at baseline and 3.5 h after intervention (i.e. exercise bout or rest). The 3.5h post-exercise biopsy was chosen based on our previous time course investigations showing robust changes in gene expression at this time point.^13,55,56,57^ Biopsy tissue was frozen and kept at -80°C until processing.

### Ethical compliance

The study complied with all ethical regulations and institutional protocols for studying human samples. All procedures associated with the research were approved by the Institutional Review Board at Ball State University and performed in accordance with relevant guidelines and regulations (IRB protocol # 123578). Subjects provided written informed consent prior to participation. Donor anonymity was preserved, and guidelines were followed regarding consent, protection of human subjects, and donor confidentiality.

### Nuclei isolation from muscle samples

Frozen human skeletal muscle samples were pulverized on dry-ice and the powder used for nuclei isolation. The protocol followed was described in Mendelev 2022 STAR protocols and Zhang 2022 Cell Reports.^58,59^ Briefly, and all on ice, RNAse inhibitor (NEB cat# MO314L) was added to the homogenization buffer (0.32 M sucrose, 1 mM EDTA, 10 mM Tris-HCl, pH 7.4, 5mM CaCl_2_, 3mM Mg(Ac)_2_, 0.1% IGEPAL CA-630), 50% OptiPrep (Stock is 60% Media from StemCell cat# 07820), 35% OptiPrep and 30% OptiPrep right before isolation. Each sample was homogenized in a Dounce glass homogenizer (1ml, VWR cat# 71000-514), and the homogenate filtered through a 40 mm cell strainer. An equal volume of 50% OptiPrep was added, and the gradient centrifuged (SW41 rotor at 17,792xg; 4C; 25min). Nuclei were collected from the interphase, washed, resuspended in 1X nuclei dilution buffer for sn multiome (10x Genomics) and counted (Cellometer) before proceeding for sn multiome assay.

### Sn multiome assay

Sn multiome was performed following the Chromium Single Cell Multiome ATAC and Gene Expression Reagent Kits V1 User Guide (10x Genomics, Pleasanton, CA). Nuclei were counted (Cellometer K2 counter), transposition was performed in 10 μl at 37°C for 60 min targeting up to 10,000 nuclei, before loading of the Chromium Chip J (PN-2000264) for GEM generation and barcoding. Samples that did not meet a minimum of 2,000 nuclei/ul at counting prior to transposition were processed in technical duplicates from the same initial pool of nuclei to obtain enough nuclei for analysis. A minimum of 1,000 total nuclei per sample were run through transposition. Following post-GEM cleanup, libraries were pre-amplified by PCR, after which the sample was split into three parts: one part for generating the snRNA-seq library, one part for the snATAC-seq library, and the rest was kept at -20°C. SnATAC and snRNA libraries were indexed for multiplexing (Chromium i7 Sample Index N, Set A kit PN-3000262, and Chromium i7 Sample Index TT, Set A kit PN-3000431 respectively). All libraries were individually checked by Bioanalyzer (Agilent) and measured by Qubit (Thermofisher). Then, libraries were pooled by ome type (snRNA or snATAC) and their quality assessed by miSeq sequencing for adjustment prior to deep sequencing using an Illumina Novaseq.

### Single-nucleus multiome primary processing and QC

CellRanger ARC 2.0.0 pipeline was run on the samples following 10x Genomics guidelines. Technical replicates of the same sample were aggregated. Seurat version 4^60–62^ and Signac version^63^ were used to perform QC on the data (Table S2, Table S3). Apoptotic, low-count cells, and doublets were identified and excluded from downstream analysis. Both apoptotic and low-count cells were identified as having lower transcript counts (≤ 500) or low number of fragments overlapping ATAC peaks (≤ 1,000). In addition, apoptotic cells had a large proportion of mitochondrial gene transcripts (≥ 5%). Doublets, on the other hand, were identified as having higher counts in either RNA (≥ 5,000-7,000 depending on sample) and ATAC (≥ 20,000). Cells with low-quality ATAC reads – defined as low percentage of reads in peaks (≤ 5%), high blacklist ratio (≥ 0.05), high nucleosome signal (≥ 10) or low TSS enrichment (≤ 2) – were also excluded.

The clustering analysis was performed using Seurat’s weighted shared nearest-neighbor graph approach.^60^ This method identifies, for each cell, its nearest neighbors based on a weighted combination of the two modalities (gene expression and chromatin accessibility). The weighted nearest-neighbor graph was then used to obtain the UMAP projection of the data. The gene expression modality was used to identify cluster cell types after determination of top markers for each cluster (Figure S1). Lymphocytes were further sub-clustered into T cells and NK cells, while phagocytes were sub-clustered into macrophages and dendritic cells.

### Differential analysis

Differential analysis was performed at both single-cell and pseudobulk levels and differentially accessible regions and differentially expressed genes were identified in a multi-step approach. Seurat (logistic regression) was used for sc differential analysis,^60–62^ while controlling for sequencing depth on the ATAC side using the peak region fragments metric. Pseudobulk count matrices were generated for both RNA and ATAC by summing gene expression and chromatin accessibility across each sample for a given cell-type. DESeq2 (Wald test) was used for pseudobulk differential analysis.^64^ All RNA analyses also included the subject id as a covariate. The following steps were carried out:

1. SC differential analysis identified features that differed between pre-vs. post-intervention samples in the exercise group with FDR-corrected p-value <= 0.05 and percent expression >= 0.1 in at least one time-point.
2. Pseudobulk differential analysis identified features that differed between pre-vs. post-intervention samples in the exercise group with p-value <= 0.05 and |log2FoldChange| > 0.3.
3. SC differential analysis identified features that differed between pre-vs. post-intervention samples in the circadian control group with FDR-corrected p-value <= 0.05 and percent expression >= 0.1 in at least one time-point.
4. Pseudobulk differential analysis identified features that differed between pre-vs. post-intervention samples in the circadian control group. Due to the small sample size of the circadian control group, p-values were not used to select features as circadian-regulated. Instead, features with a difference between pseudobulk exercise |log2FoldChange| and pseudobulk control |log2FoldChange| of less than 0.1 were identified as circadian control differential features.

The final set of exercise-regulated features includes features that are differentially regulated in the exercise group defined as the intersection of the sc analysis (step 1) and pseudobulk analysis (step 2). Circadian control features, defined as the union of the sc analysis (step 3) and pseudobulk analysis (step 4), are excluded from the final set of exercise-related features.

### Pathway Enrichment

ClusterProfiler version 4 was used to find enriched pathways for differential genes, as defined above.^65,66^ The biological process ontology is used for enrichment and the background genes are the sc cell-type markers. Figure 2C and Figure S4 show the consolidated top ten terms per cell type, sorted by LFC.

### Curating a Previously Reported Exercise-Response Geneset

The previously reported exercise-response geneset was downloaded from the GitHub repository associated with the meta-analysis by Amar et al.^23^ The downloaded data was filtered to only include genes from acute muscle exercise-response studies with a meta-analysis p-value of < 0.05.

### PLIER Latent Variable Analysis

A pseudobulk count matrix was generated for RNA by averaging normalized gene expression across each sample for a given cell-type. To generate a pseudobulk matrix for ATAC-seq, each chromatin region was first annotated as a promoter region^67^ for a gene if it was located within 3kb upstream or downstream of the gene transcription start site (TSS). A cumulative accessibility value was computed for each gene in each cell by summing the region accessibilities for all promoter regions. Finally, a pseudobulk count matrix for the ATAC-seq was calculated by averaging the total promoter region accessibility for each gene across each sample for a given cell-type.

The fiber-type RNA-seq and ATAC-seq pseudobulk profiles (fast, slow and intermediate) were z-scored by feature for RNA-seq and ATAC-seq separately. Pathway-Level Information ExtractoR (PLIER)^38^ was applied to the concatenated gene-indexed RNA-seq and ATAC-seq matrices, identifying 12 latent variables of shared measurement patterns across samples, cell types, and omes. The statistical significance of the response to exercise (LV value) was measured within each ome separately by a paired t-test and adjusted for multiple hypotheses by the Holm-Bonferroni method. Associated pathways were identified for the LVs as positive U matrix values in PLIER.

### MAGICAL Analysis

Multiome Accessibility Gene Integration Calling And Looping (MAGICAL) was applied to the sc multiome data to identify regulatory circuits that include transcription factors, associated chromatin sites and target genes.^40^ MAGICAL is a Bayesian framework that iteratively models chromatin accessibility and gene expression variation across cells and samples in each cell type. It estimates the confidence of TF binding at open chromatin regions and also their linkages to target genes as regulatory circuits. MAGICAL built candidate exercise-associated circuits by mapping TFs to differentially accessible regions (DAR) within each cell type using motif sequences from the chromVARmotifs library.^68^ Then, it linked the binding sites to DEGs in that cell type by genomic localization within +/-500Kb. Here, DAR and DEG were selected if they were differential at either sc or pseudobulk level per cell type (see Differential Analysis).

### CREMA Analysis

Control of Regulation Extracted from Multiomics Assays (CREMA) is a method for regulatory circuit inference in multiome data that scans for potential TF binding sites by motif analysis in the +/-100kb region around the transcription start site in a peak-agnostic manner, and uses a linear model to find regulatory circuits supported by the coincidence of the expression of the target gene, the expression of the TF and the local chromatin accessibility of the TF’s binding site.^39^ Samples for each subject (both pre and post timepoints) were pooled, and CREMA was applied to extract regulatory circuits specific to each subject and cell type. Circuits were selected if they were replicated in at least two of the four subjects in the exercise group and satisfied at least one of the following two criteria: 1) the target gene and the TF both showed significant differential expression between pre- and post-exercise; 2) the target gene and the *cis*-regulatory site both showed significant differential expression and differential accessibility respectively between pre- and post-exercise. Here, differential expression and accessibility were defined as being differential at either sc or pseudobulk level per cell type (see Differential Analysis). Because this analysis focused on regulatory circuits with positive regulation, the fold-change was required to be in the same direction.

### Integrative Analysis to Define Regulatory Circuits

The regulatory circuits extracted by MAGICAL and CREMA were integrated by taking the union of regulatory circuits as defined by the two methods. Specifically, for each target gene, all the TFs and *cis*-regulatory regions found by the two methods were aggregated as the regulatory circuits for this target gene.

### Data availability

The datasets generated in the present study are deposited in GEO (accession ID: GSE24006).

### Code availability

Any computational code used in the paper is available upon request.

### Material availability

This study did not generate new unique reagents.

### Supplemental Information Titles and Legends

**Figure S1:** Single-cell primary processing: feature plots showing marker genes or QC metrics on the UMAP projection. Marker genes include *MYH2* (slow/intermediate), *MYH7* (fast/intermediate), *VWF* (endothelial cells), *RGS5* (pericytes), *FBN1* (FBN1+ FAP cells), LUM (LUM+ FAP cells), *MAF* (macrophages), *HLA-DRA* (dendritic cells), *PAX7* (satellite cells), *CD3D* (T cells), *NKG7* (NK cells), *MS4A1* (B cells), *KIT* (mast cells). QC metrics include nCount_RNA (the number of RNA UMIs per cell), nCount_ATAC (the number of ATAC UMIs per cell), and percent.mt (percentage of mitochondrial reads per cell).

**Figure S2:** Cell-type proportions are roughly comparable between pre and post time-points. (A) Boxplots reveal frequency of each cell-type within each sample and are split by time-point and group (n = 2 control samples per time-point and n = 4 exercise samples per time-point). Colors denote cell-type with a darker tint indicating the post time-point. To enable more consistent comparison, fibers are excluded during computation of cell-type frequency. Change between pre and post cell-type proportions is assessed via Holm-Bonferroni adjusted paired t-test with ns: p > 0.05, *: p <= 0.05, **: p <= 0.01, ***: p <= 0.001, ****: p <= 0.0001 (B) Barplot indicates frequency of each cell-type, including fibers, for each sample. Subjects Ctrl-1, Ex-1, and Ex-2 are female; subjects Ctrl-2, Ex-3, and Ex-4 are male.

**Figure S3:** Differential analysis of circadian controls and overlaps of cell-type differential features. (A-B) Volcano plot showing differentially expressed genes pre vs. post (computed at single-cell resolution) for slow fibers in the exercise group (A) and control group (B). *FKBP5* is labeled in both panels and is highly significant with a log_2_ fold change < -2. (C-D) Volcano plots show the distributions of upregulated and downregulated genes (C) and chromatin regions (D) for each cell-type in circadian controls. These are the features that were excluded from the set of exercise-response features prior to downstream analysis. (E-F) UpSet plots of DEG (E) and DAR (F). Cell-types with fewer than 10 differential features and intersections with 1 differential feature are not shown.

**Figure S4:** Enrichment analysis for all cell-types. Enrichment analysis of biological processes in each cell-type shows functionally different transcriptome profiles for each cell-type. Color represents the percentage of upregulated genes in a given pathway and larger dot radius denotes more significant biological function.

**Figure S5:** PLIER Supplementary Panels. (A) Heatmap represents summary values for all PLIER-identified LVs. Each column represents one fiber-type sample; columns are ordered by ome type, exercise group, and time-point. Each row represents one LV. (B) Heatmap of enriched pathways associated with select latent variables. Box color reflects AUC for association of pathway to LV.

**Supplementary Table S1**: Subject Characteristics

**Supplementary Table S2:** QC RNA-seq Metrics

**Supplementary Table S3:** QC ATAC-seq Metrics

**Supplementary Table S4:** PPAR𝜹 Targets

