## Supplemental Information for "Integrated single-cell multiome analysis reveals muscle fiber-type gene regulatory circuitry modulated by endurance exercise"

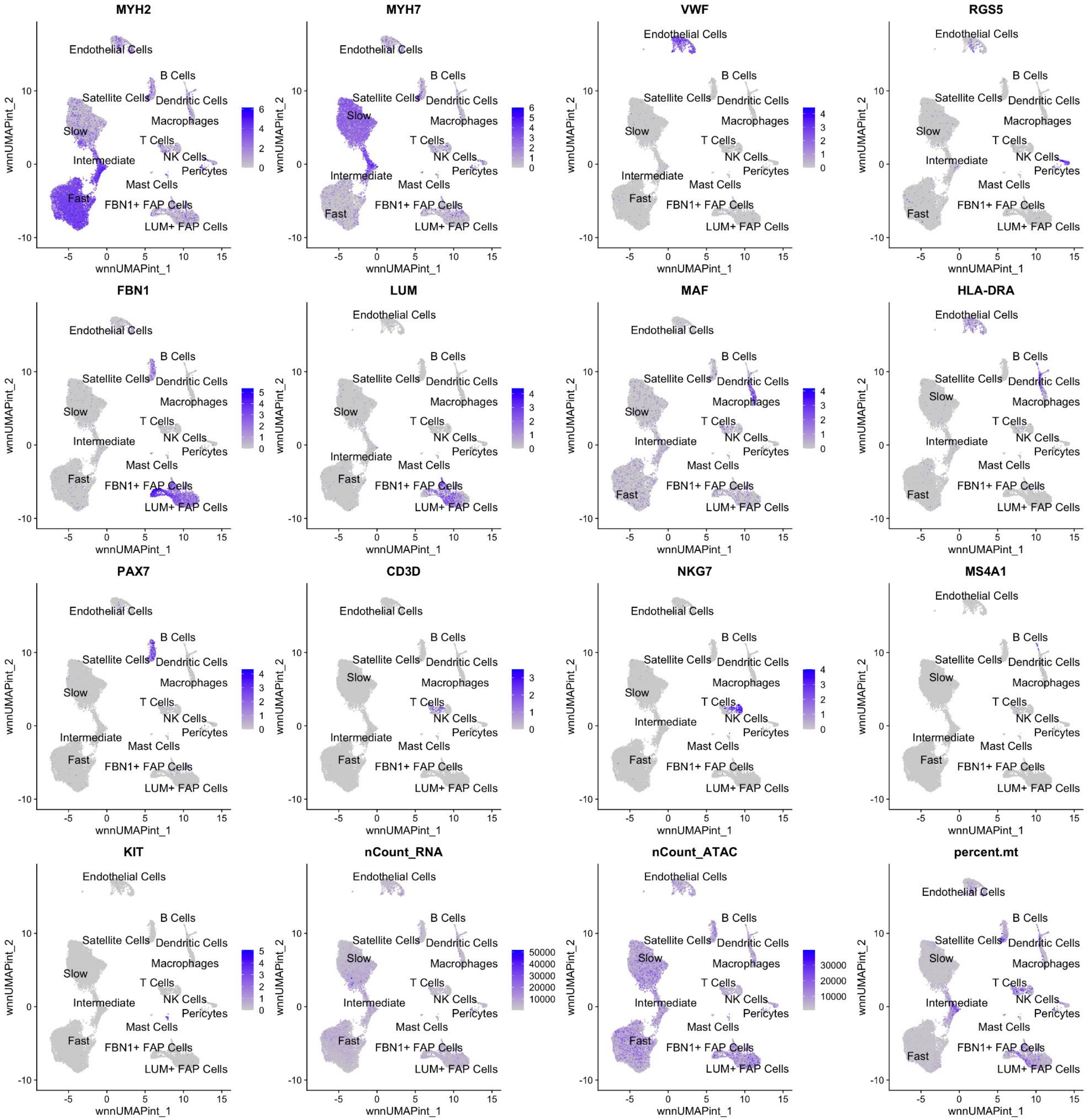


**Figure S1**: Single-cell primary processing: feature plots showing marker genes or QC metrics on the UMAP projection. Marker genes include *MYH2* (slow/intermediate), *MYH7* (fast/intermediate), *VWF* (endothelial cells), *RGS5* (pericytes), *FBN1* (FBN1+ FAP cells), LUM (LUM+ FAP cells), *MAF* (macrophages), *HLA-DRA* (dendritic cells), *PAX7* (satellite cells), *CD3D* (T cells), *NKG7* (NK cells), *MS4A1* (B cells), *KIT* (mast cells). QC metrics include nCount_RNA (the number of RNA UMIs per cell), nCount_ATAC (the number of ATAC UMIs per cell), and percent.mt (percentage of mitochondrial reads per cell).

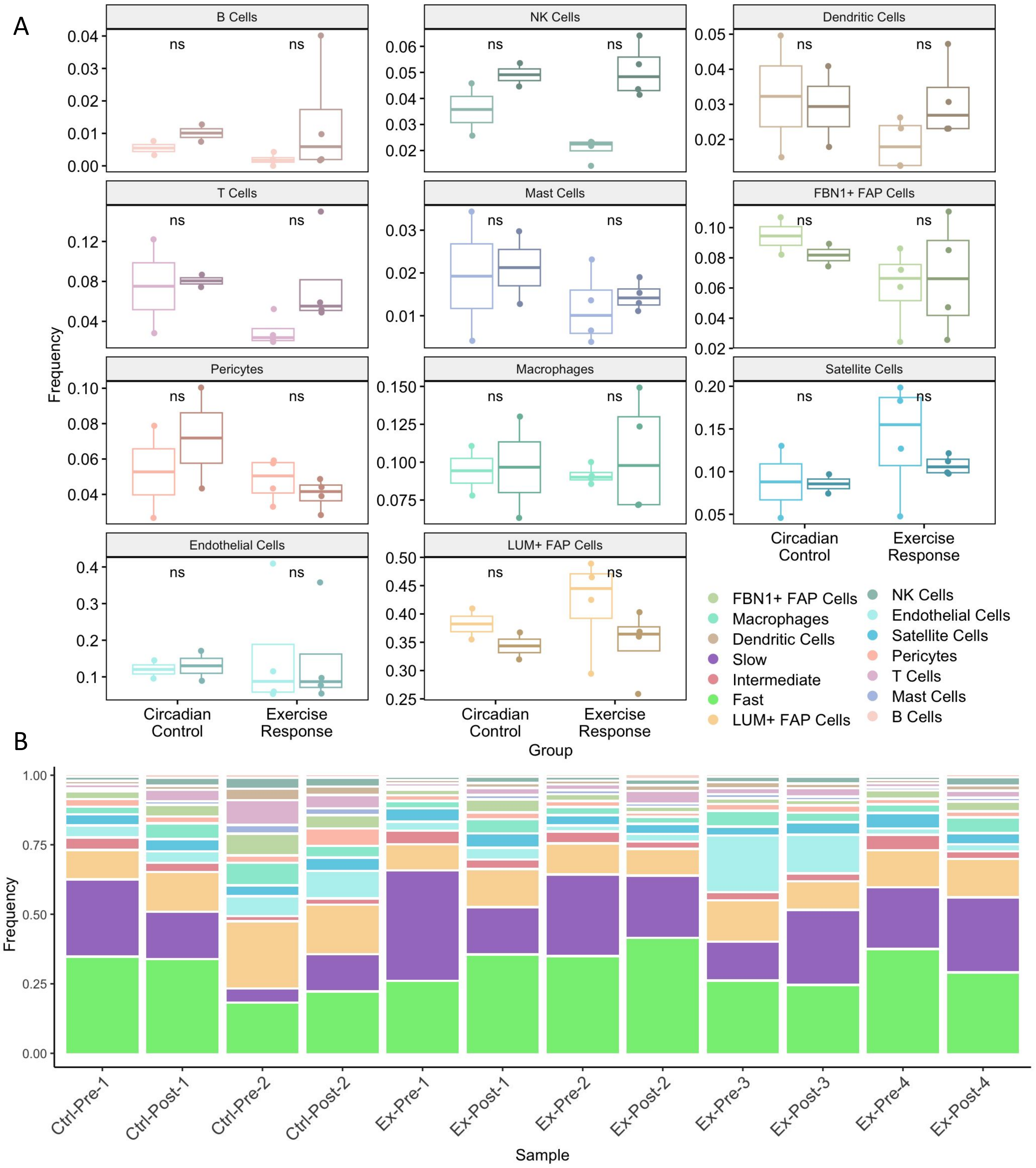


**Figure S2**: Cell-type proportions are roughly comparable between pre and post time-points. (A) Boxplots reveal frequency of each cell-type within each sample and are split by time-point and group (n = 2 control samples per time-point and n = 4 exercise samples per time-point). Colors denote cell-type with a darker tint indicating the post time-point. To enable more consistent comparison, fibers are excluded during computation of cell-type frequency. Change between pre and post cell-type proportions is assessed via Holm-Bonferroni adjusted paired t-test with ns: $p > 0.05$, *: $p <= 0.05$, **: $p <= 0.01$, ***: $p <= 0.001$, ****: $p <= 0.0001$ (B) Barplot indicates frequency of each cell-type, including fibers, for each sample. Subjects Ctrl-1, Ex-1, and Ex-2 are female; subjects Ctrl-2, Ex-3, and Ex-4 are male.

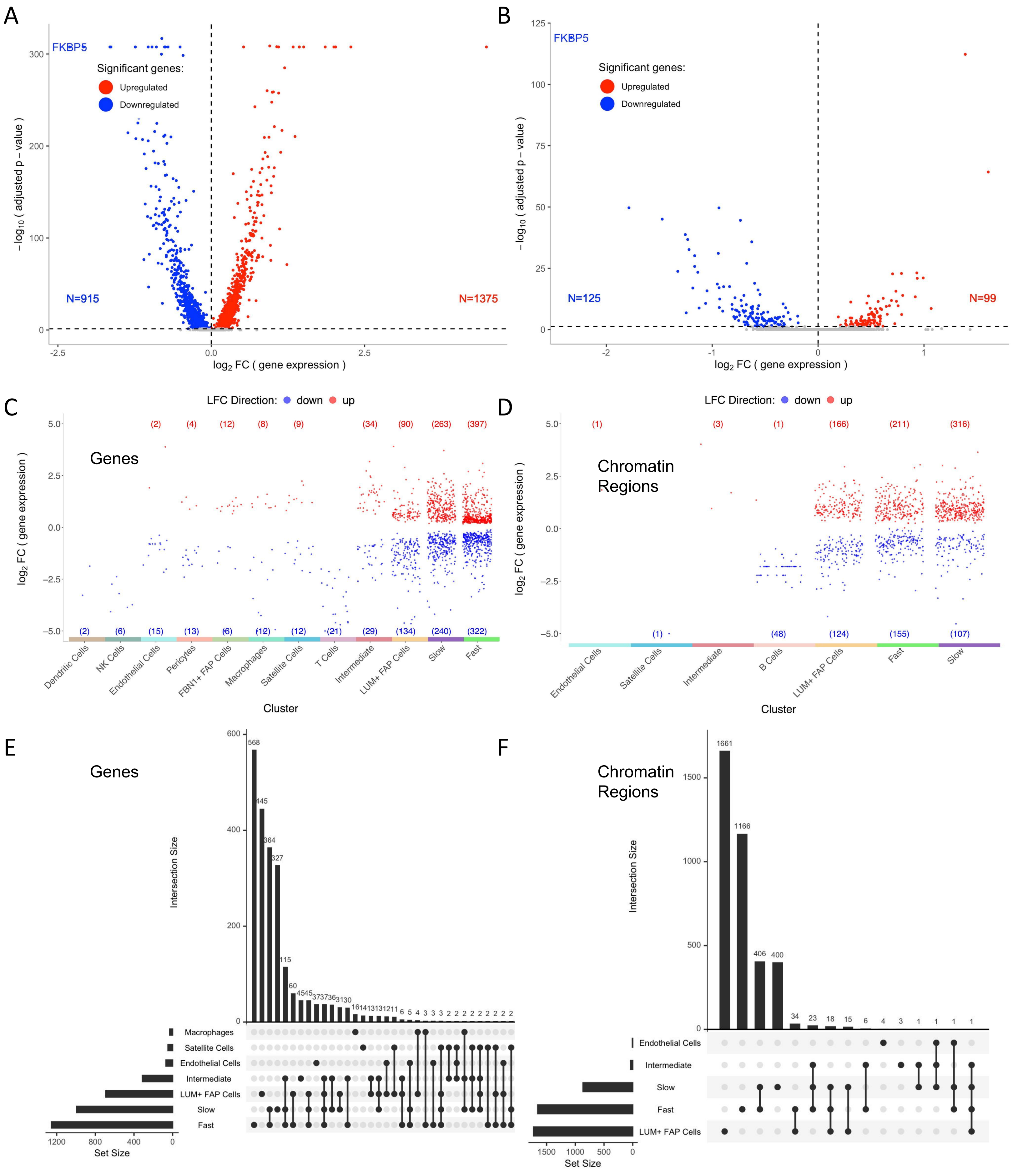


**Figure S3**: Differential analysis of circadian controls and overlaps of cell-type differential features. (A-B) Volcano plot showing differentially expressed genes pre vs. post (computed at single-cell resolution) for slow fibers in the exercise group (A) and control group (B). *FKBP5* is labeled in both panels and is highly significant with a $\log_2$ fold change < -2. (C-D) Volcano plots show the distributions of upregulated and downregulated genes (C) and chromatin regions (D) for each cell-type in circadian controls. These are the features that were excluded from the set of exercise-response features prior to downstream analysis. (E-F) UpSet plots of DEG (E) and DAR (F). Cell-types with fewer than 10 differential features and intersections with 1 differential feature are not shown.

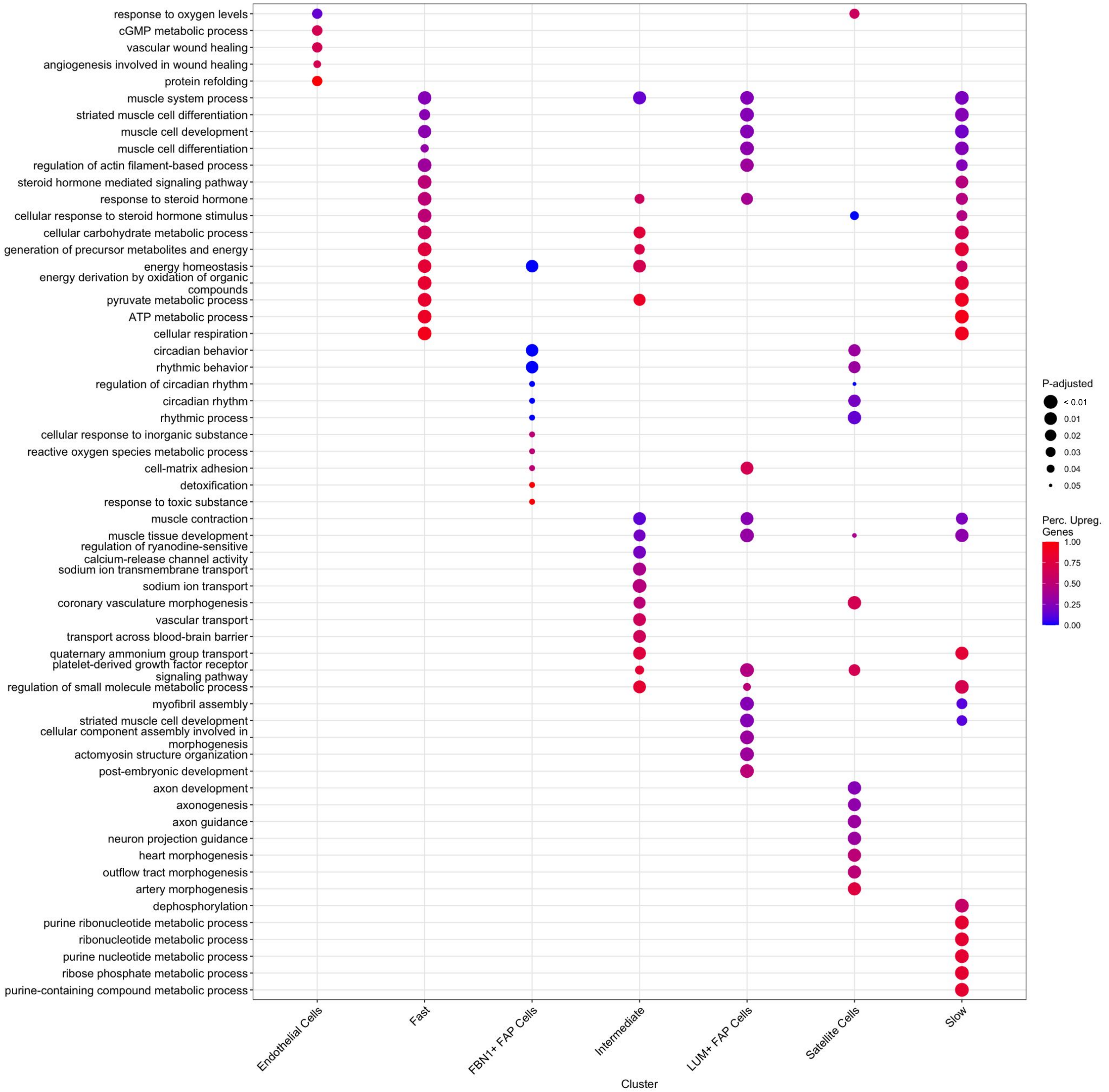


**Figure S4**: Enrichment analysis for all cell-types. Enrichment analysis of biological processes in each cell-type shows functionally different transcriptome profiles for each cell-type. Color represents the percentage of upregulated genes in a given pathway and larger dot radius denotes more significant biological function.

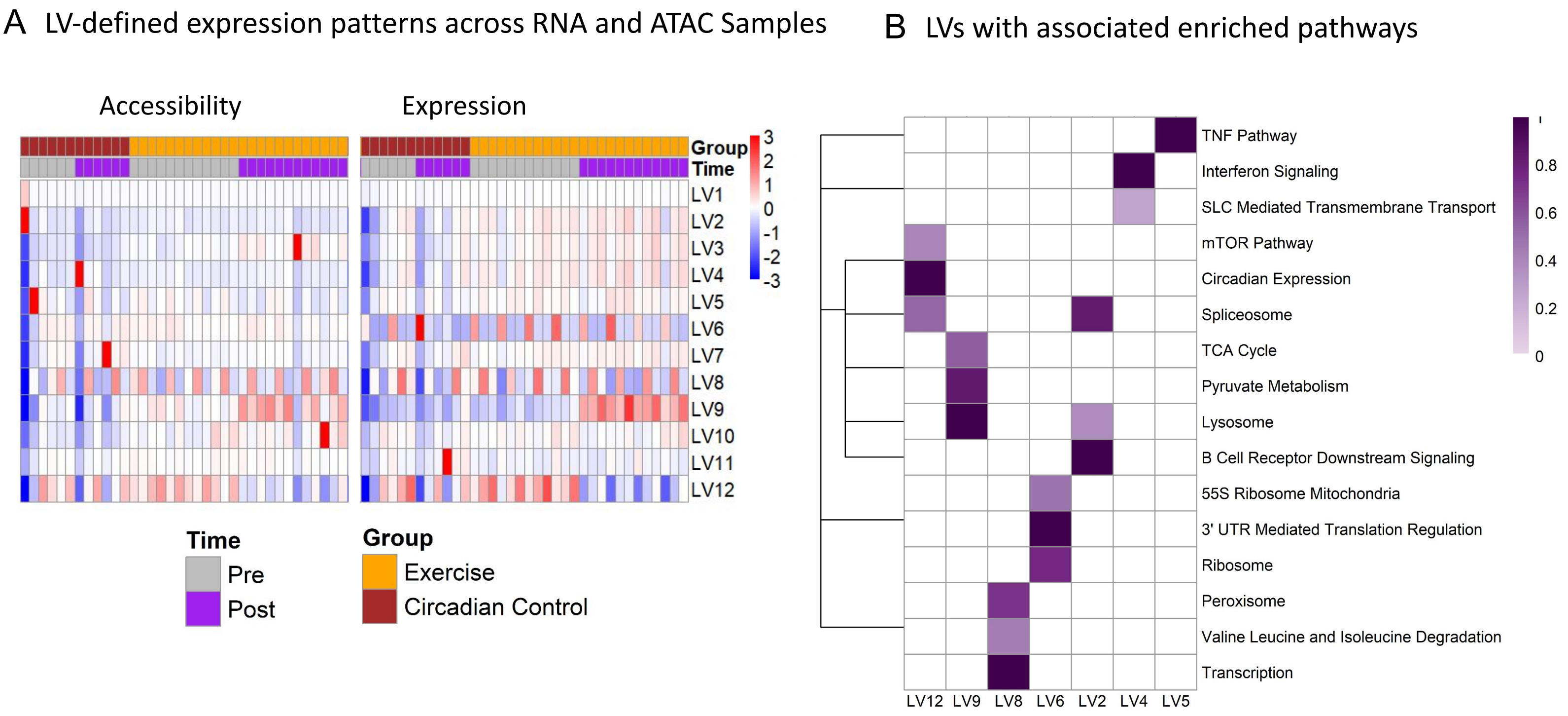


**Figure S5**: PLIER Supplementary Panels. (A) Heatmap represents summary values for all PLIER-identified LVs. Each column represents one fiber-type sample; columns are ordered by ome type, exercise group, and time-point. Each row represents one LV. (B) Heatmap of enriched pathways associated with select latent variables. Box color reflects AUC for association of pathway to LV.

**Supplementary Table S1**: Subject Characteristics

| Subject | Sex | Group | Age | Height (cm) | Weight (kg) | BMI (kg·m$^2$) | Body Fat % | Absolute $VO_2$max(L/min) | Relative $VO_2$max(ml/kg/min) |
|---|---|---|---|---|---|---|---|---|---|
| **E** | F | Exercise | 23 | 173 | 72.90 | 24.4 | 34.6 | 2.86 | 39.24 |
| **G** | M | Exercise | 23 | 175 | 88.24 | 28.8 | 24.4 | 3.49 | 39.57 |
| **I** | F | Exercise | 26 | 164.2 | 68.9 | 25.6 | 30.8 | 2.78 | 40.37 |
| **J** | M | Exercise | 26 | 175 | 91.3 | 29.8 | 33.1 | 4.31 | 47.24 |
| **L** | M | Control | 23 | 184 | 82.7 | 24.4 | 19.4 | 3.89 | 47.02 |
| **N** | F | Control | 24 | 151 | 59.63 | 26.2 | 34.6 | 3.03 | 50.76 |

BMI, body mass index; F, female; M, male; $VO_2$max, maximal oxygen consumption.

**Supplementary Table S2**: QC RNA-seq Metrics

| Subj | Time | Estimated number of nuclei | Mean reads/cell | Median UMI/cell | Median genes/cell | Reads mapped to transcriptome | Reads mapped to genome | Fraction of transcriptomic reads in cells |
|---|---|---|---|---|---|---|---|---|
| E | 1.0 | 7960 | 52666 | 3906 | 1727 | 0.30 | 0.93 | 0.80 |
| E | 2.0 | 5391 | 67791 | 3865 | 1841 | 0.32 | 0.93 | 0.77 |
| G | 1.0 | 3003 | 131480 | 4386 | 1950 | 0.28 | 0.92 | 0.80 |
| G | 2.0 | 4335 | 91301 | 3328 | 1660 | 0.33 | 0.92 | 0.78 |
| I | 1.0 | 5854 | 48101 | 3660 | 1652 | 0.31 | 0.94 | 0.84 |
| I | 2.0 | 4608 | 57218 | 3356 | 1678 | 0.31 | 0.94 | 0.80 |
| J | 1.0 | 8772 | 32451 | 3797 | 1780 | 0.31 | 0.94 | 0.85 |
| J | 2.0 | 2524 | 109593 | 3944 | 1874 | 0.35 | 0.94 | 0.81 |
| L | 1.0 | 632 | 667124 | 4788 | 2115 | 0.41 | 0.91 | 0.61 |
| L | 2.0 | 961 | 502041 | 4531 | 2035 | 0.49 | 0.91 | 0.49 |
| N | 1.0 | 6499 | 67568 | 4064 | 1805 | 0.34 | 0.93 | 0.72 |
| N | 2.0 | 1372 | 557108 | 5070 | 2194 | 0.29 | 0.92 | 0.71 |

**Supplementary Table S3**: QC ATAC-seq Metrics

| Subj | Time | Median high-quality fragments/cell | Fraction of transposition events in peaks in cells | Number of peaks | Fraction of genome in peaks | TSS enrichment score | Confidently mapped read pairs | Fraction of high quality fragments overlapping peaks |
|---|---|---|---|---|---|---|---|---|
| E | 1 | 18055 | 0.332 | 84518 | 0.023 | 5.47 | 0.93 | 0.33 |
| E | 2 | 20117 | 0.320 | 78916 | 0.022 | 5.02 | 0.93 | 0.32 |
| G | 1 | 25973 | 0.378 | 97398 | 0.027 | 6.09 | 0.92 | 0.38 |
| G | 2 | 18539 | 0.314 | 75830 | 0.021 | 4.74 | 0.92 | 0.31 |
| I | 1 | 10428.5 | 0.347 | 92023 | 0.026 | 6.27 | 0.94 | 0.38 |
| I | 2 | 8209 | 0.188 | 54498 | 0.015 | 4.24 | 0.93 | 0.21 |
| J | 1 | 10601 | 0.348 | 100678 | 0.028 | 6.05 | 0.94 | 0.38 |
| J | 2 | 25056 | 0.314 | 83645 | 0.023 | 5.16 | 0.93 | 0.34 |
| L | 1 | 26888.5 | 0.208 | 35652 | 0.010 | 3.77 | 0.91 | 0.21 |
| L | 2 | 21707 | 0.156 | 29919 | 0.008 | 3.28 | 0.91 | 0.16 |
| N | 1 | 10489 | 0.378 | 92724 | 0.026 | 6.50 | 0.93 | 0.38 |
| N | 2 | 23827 | 0.344 | 66074 | 0.018 | 5.09 | 0.92 | 0.34 |

**Supplementary Table S4**: PPAR$\delta$ Targets

| **Cell type** | **PPAR$\delta$ targets** |
|---|---|
| Fast fiber | ADORA2A-AS1, P4HA2-AS1, DPP6, C16orf46, TBC1D1, PTTG2, FABP3, PTPRG, CCDC197, RELL1, SERPINB9, ACKR3, SLC25A18, SLC26A9, VEGFA, CASTOR2, SPIRE1, IQCC, RNF144B, TFEB, SERINC2, ANGPTL2, MBP, CDHR3, SLC41A1, SLC38A4, RFTN1, RAB29, ITIH6, RAB7B, RHBDF1, MOCS1, MIA2, PHKG1, STC2, PTPRJ, SLC30A2, FAM83E, PDE11A, CDS2, ADM, OLFM1, RTN4RL1, LANCL1-AS1, NDRG1, GCOM1, CDKN1A, SLC22A23, SLC22A5, TMEM52, HADHB, LARGE-AS1, CES3, SPTBN4, RSPH9, C3orf35, CREB3L2, MID1IP1, SH3BP2, EML1, ZNF366, PSMG4, CPEB4, C4orf19, FAM43A, VGLL2, TMEM164, MIR3936HG, UQCRFS1, LIMD1-AS1, RIPOR1, DAAM2, LARGE1, RRS1, HHATL, POR, ZBTB43, ABCC1, ANKRD9, ERAP1, TSPAN9, PARVA, KLHL40, PSMD8, RANBP10, IGF2R, TMEM140, SCARF1, TRAF5, TECPR2, PPTC7, SLC25A5, DHDDS, KCNC1, LINC01968, SLC16A1-AS1, CALML6, IAH1, SOCS7, FARP1, RAB11FIP3, GANC, SAMD4A, HIVEP2, SLC22A4, ASAP3, DNAJA4, DLGAP4-AS1, IBTK, ZCCHC17, EIF4EBP2, ZMIZ1, NAGLU, SFMBT1, TNS3, CCDC13, CLIP4, KIF13A, MPC1, NCR3LG1, COLQ, RHOBTB2, DLGAP4, RAD9B, C19orf12, EPS15L1, TKT, UBE2O, ABHD3, PTCD3, GOLGA4, RBMS3-AS3, CRK, AFG3L2, LINC01909, RGS3, OSBPL2, MIR1-1HG, NODAL, RBM38, ZNF385C, PRKCD, LGALSL, PDCD11, SLC22A1, DEK, GBF1, ADHFE1, ABCC8, PIM1, PRAG1, ARMH3, TOGARAM2, IDI2-AS1, SPECC1L, FKBP9, SEPTIN11, CD79A, ZBED6, SH3KBP1, GPR179, LCA5L, PARM1, LINC01754, PALMD, RETSAT, WDR60, CHD2, STAT3, SETD3, ITGB1BP1, SLC37A1, PITPNM2, ACSBG1, BCL2L14, PREB, PAFAH2, PARVB, SLC11A2, SGSM1, C16orf70, PTPN3, MTHFD1, ACACB, SLC4A4, MAN1C1, FNBP1L, ATP5MG, RILPL1, DUS2, NCOA3, GREB1L, JMJD6, UBAC1, ZNF609, FAM181A, ETV6, SLC31A1, LRIG2-DT, PHF20, DPEP2, CES2, PLAGL1, CLIC5, SLC29A1, BCAT2, ATP5MD, MTCH2, CEP68, GLCCI1, SNHG1, LACTB, ABHD2, CIAO2A, CFL2, CAMSAP1, RHBDF2, MIOS, CCDC28B, ZCWPW2, ANKRD28, HSPB1, DHRS7C, SELENON, PLEKHA4, SLC66A2, WDR37, PFKFB1, CGREF1, ABHD1, ARHGEF10L, GEM, MIPEP, ANP32A, STK24, GPR157, RDH5, SH2B3, TBC1D22A, PDCD10, GREB1, PPP4C |
| Slow fiber | CGREF1, KLF13, CNNM1, TKT, GCOM1, JUNB, LINC00598, AMOTL1, IDH3A, ACSBG1, PREB, TRIM63, GDAP1L1, NRP1, STAT3, CCM2, ZMIZ1, ACOT11, MAP4K3-DT, SLC41A1, UBAC1, FAM151A, NOP16, PKIG, ZNF44, ATP5MG, EPDR1, GOT1, HOOK2 |
